# Effects of medial septum low-frequency deep brain stimulation on pentylenetetrazol-induced seizures in rats

**DOI:** 10.64898/2026.04.24.720729

**Authors:** Alejandra Garay-Cortes, Salvador Almazan-Alvarado, Victor Manuel Magdaleno-Madrigal, Hiram Luna-Munguia

## Abstract

**Background:** Invasive neuromodulation may be used in patients if seizure medications fail and surgery is not an option. However, moderate success is achieved and improved paradigms are required. The medial septum has been considered a suitable target for the treatment of temporal lobe epilepsy due to its location and connectivity.

**Objective:** To assess the effect of medial septum low-frequency deep brain stimulation to inhibit pentylenetetrazole (PTZ)-induced seizures.

**Methods:** Male Sprague-Dawley rats were stereotaxically implanted in the medial septum and left dorsal hippocampus one week prior to the experimental protocols. Then, the animals were assigned to three experimental groups: 1) 10 Hz + PTZ (n=3); 2) 5 Hz + PTZ (n=7); and 3) 5 Hz (n=7). The stimulation consisted of a 30 min train of biphasic square-wave pulses at a current of 150 µA and a pulse duration of 1 ms. Rats were subjected to the experimental protocol every 24 h for seven consecutive days.

**Results:** Subjects exposed to the 10 Hz died after the first PTZ injection. The 5 Hz stimulation not only prevented the animals’ death, but also induced a protective effect against generalization. Surprisingly, in both 5 Hz groups, septal and hippocampal spike-wave-like discharges were detected (mainly integrated by theta oscillations). This phenomenon was correlated with the generalization avoidance.

**Conclusions:** While this study is preclinical in nature, our findings underscore the potential of using low-frequency medial septum stimulation for future clinical applications.

**Graphical Abstract:** 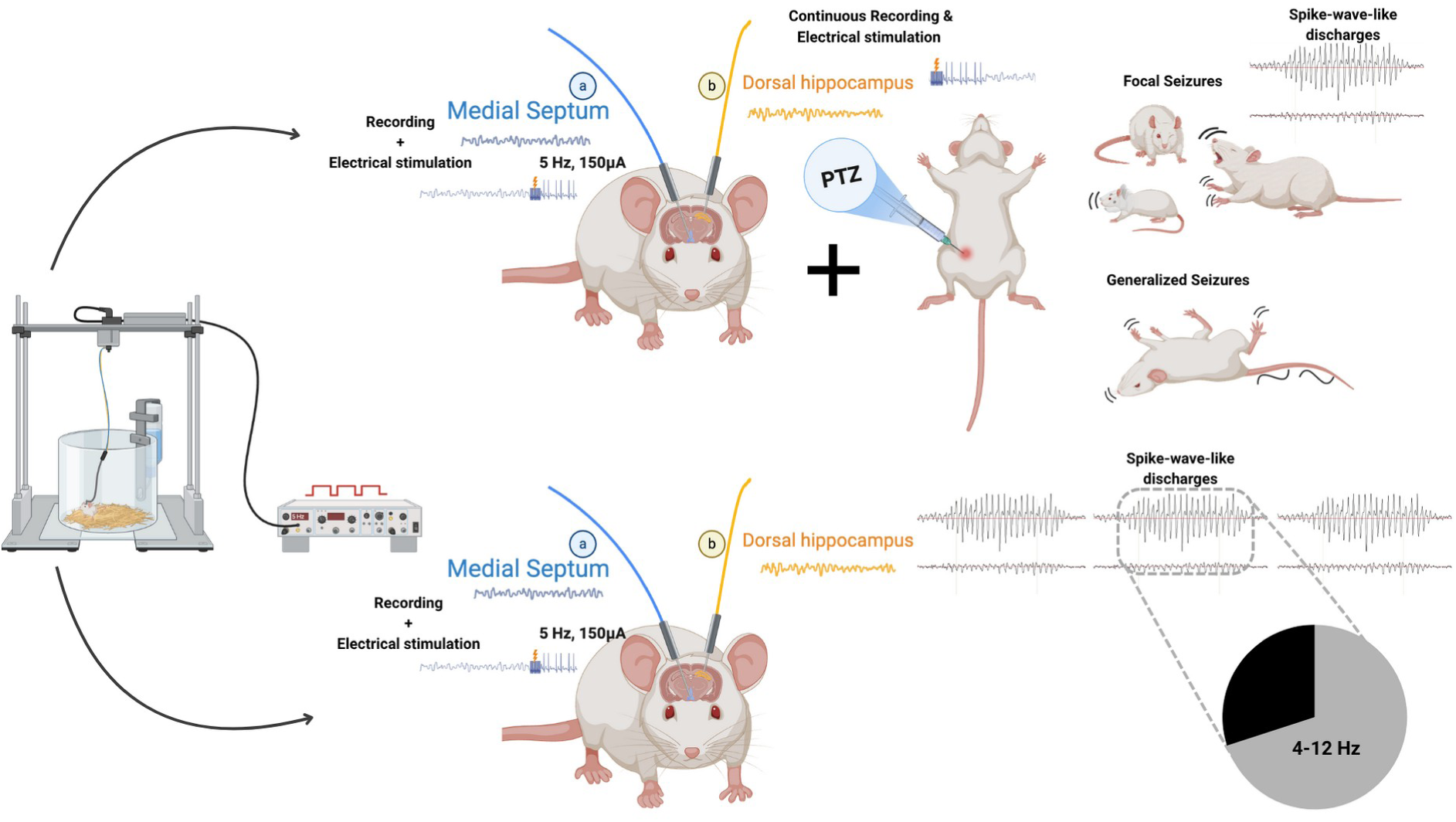

## 1. INTRODUCTION

Over the last three decades, neuromodulation has been considered as an alternative therapeutic option for patients with drug-resistant epilepsy who are not candidates for surgical resection of the epileptogenic area [1–3]. In the case of deep brain stimulation of the anterior nucleus of the thalamus, the Food and Drug Administration (FDA) has reported it as an effective treatment for refractory patients, achieving seizure freedom or seizure reduction for several months [4,5]. The effectiveness depends on the type of epilepsy and the stimulated brain region [6,7], as well as the applied stimulation pattern (frequency, current intensity, waveform shapes) [8–10]. To date, the physiological mechanisms involved in seizure management have not been fully elucidated. *In vitro* research and experimental animal studies suggest that the mechanisms may include: 1) an increase in extracellular potassium concentration and neuronal depolarization block during stimulation [11,12]; 2) a prevention or delay of the synchronization and propagation of paroxysmal activity between cortical and subcortical regions [13]; and 3) an augmented release of gamma-aminobutyric acid (GABA) into the extracellular space of the stimulated area [14,15]. Improving this treatment modality has been complicated, as there is a limited understanding of the mechanisms underlying the antiseizure effects of deep brain stimulation and how seizures engage specific neuronal circuitry.

The search for new cerebral stimulation targets is necessary to clinically provide alternatives for better seizure control. The medial septum, a small gray matter nucleus located in the middle of the basal forebrain, has recently been proposed as a field of interest [16,17]. In rodents, this structure contains interconnected cholinergic, GABAergic, and glutamatergic neurons [18–21], and it is the primary pacemaker of the hippocampal theta rhythm (4-12 Hz) [22–26]. Multiple studies in animal models of temporal lobe epilepsy have described a disruption of hippocampal theta oscillations [27–34]. Recent studies, using electrical and optogenetic stimulation in the medial septum have reported that increased theta rhythms are correlated with periods of reduced epileptiform activity [35–41]. The aim of this study is to provide evidence supporting the idea that theta rhythm restoration via low-frequency deep brain stimulation of the medial septum could be an effective neuromodulation treatment for seizure control. For this purpose, we used the pentylenetetrazole (PTZ) screening test, a widely used animal seizure model employed in the search for new antiseizure drugs [42–44].

## 2. METHODS

### 2.1 Animals

The procedures were performed in accordance with the Mexican Federal regulations for animal experimentation (NOM-602-ZOO-1999) and the ARRIVE guidelines [45]. The experimental design was approved by the Institute of Neurobiology Animal Care and Ethics Committee (protocol 105A).

All subjects included in the study were age-matched (3 months, 280-320 g body weight) male Sprague-Dawley rats provided by the Institute’s animal facility. The rats were housed in a room under controlled illumination (12-h light/dark cycle) and environmental conditions (20-22°C, 50-60% humidity). All had free access to food and water and were acclimatized to the room conditions for at least one week before any experimental manipulation. Efforts were made to minimize the number of animals used.

2.2 *In vivo* magnetic resonance imaging

The acquisition protocol was carried out four days before the surgery (see below) at the National Laboratory for magnetic resonance imaging (MRI) using a 7 T scanner (Bruker Pharmascan 70/16US). *In vivo* T2 images were acquired using a 72 mm inner-diameter volume coil for transmission and a 2×2 rat head array coil for reception (Bruker, Ettlingen, Germany). The aim of this 5 min anatomical scan was to detect unexpected brain injuries in animals assigned to the surgical procedure [46]. Rats showing abnormalities were discarded.

### 2.3 Surgery

Seventeen healthy rats were used as experimental subjects. The top of their heads was shaved once they were anesthetized with a ketamine/xylazine mix (70/10 mg/kg ip, respectively). Then, the animals were placed in a Stoelting stereotaxic frame, and their body temperature was controlled and maintained via a heating pad. An incision was made midline along the scalp, and the skull was exposed using sterile surgical techniques. Two stainless steel screws were placed in the skull over the frontal cortex (left used as ground for the medial septum tripolar electrode; right used as a skull anchor), and one screw was placed over the right parietal cortex (used as ground for the left dorsal hippocampus bipolar electrode). The craniotomy for the medial septum local field potential recording electrode, which we constructed, was performed at the following coordinates: anteroposterior +0.5 mm, lateral −2 mm, ventral from skull surface −7 mm, 20° angle in the coronal plane. The coordinates for the left dorsal hippocampus were: anteroposterior −3.3 mm, lateral −2.2 mm, ventral from skull surface −3.4 mm. All coordinates were obtained from the Paxinos and Watson rat brain atlas [47] with bregma as the reference point (Fig. 1A and 1B). The entire implant was secured with dental cement. All animals received an intramuscular post-operative analgesic treatment (ketorolac 30 mg/ml; PiSA) every 24 h for two days, and allowed to recover for 7 days. Following surgery, the rats were individually housed in Plexiglas cages.

**Figure 1.**
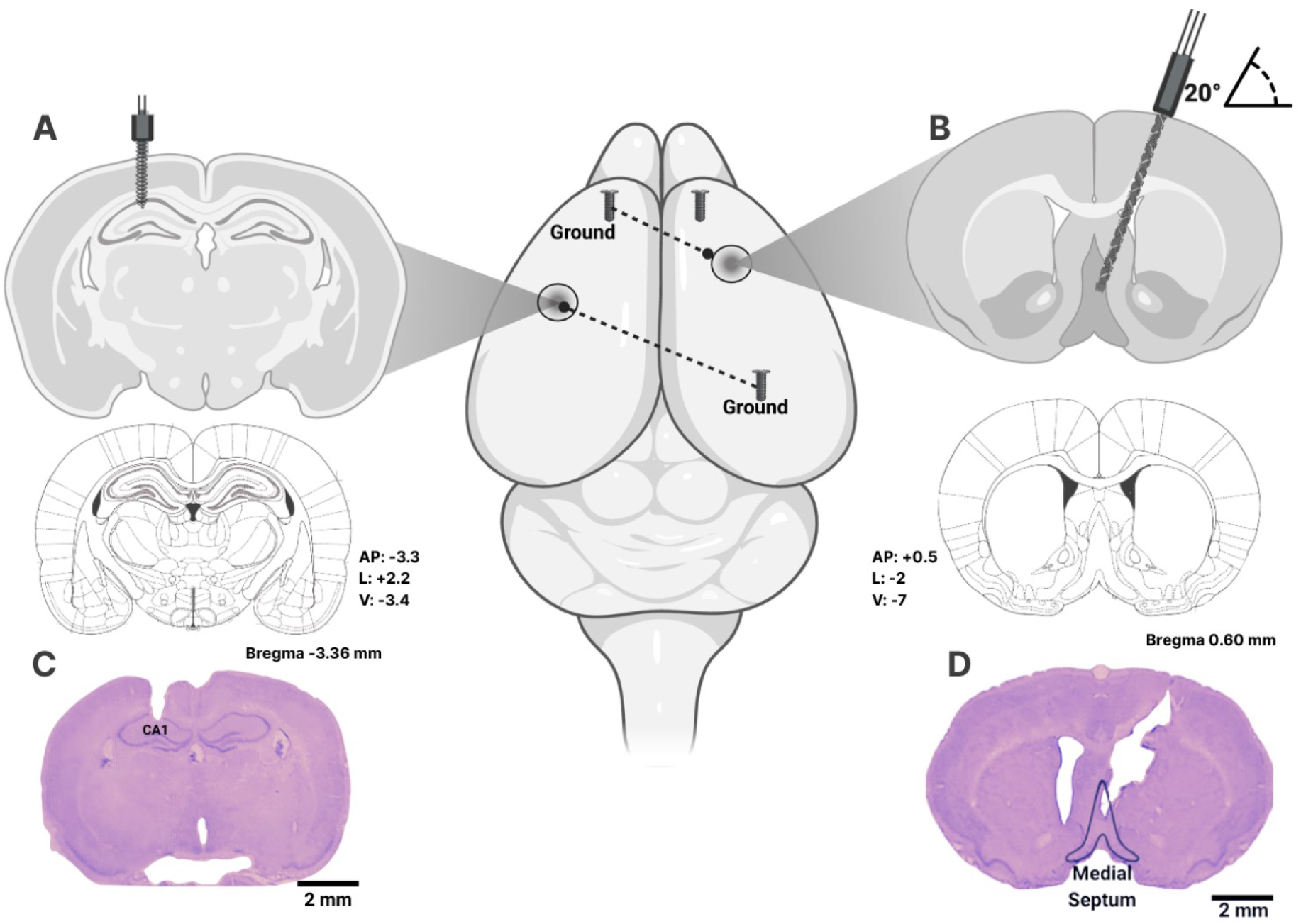
Nissl staining at the electrode placement coordinates. Stainless steel screws were placed over the frontal cortex and right parietal cortex. The craniotomy for the left dorsal hippocampus and medial septum local field potential recording electrodes was performed at specific coordinates **(A, B)**. Nissl staining showing the left dorsal hippocampus (CA1) **(C)** and medial septum tip location **(D)**. Scale bars: 2 mm.

### 2.4 Recording of local field potentials and electrical stimulation

On the day of the experiment, the animals were placed in a Plexiglas cylinder. The implanted electrodes were connected by insulated wires to a Grass 78H polysomnograph amplifier and a Grass Instruments S88 stimulator. After a 10 min adaptation period, the basal local field potential signal of the medial septum and left dorsal hippocampus was simultaneously acquired and sampled at 500 Hz for 5 min. Signals were amplified, filtered using a band pass filter between 3 and 100 Hz, and digitized at 1024 Hz with an analog-digital converter we built.

The animals were randomly assigned to three experimental groups: 1) the 10 Hz + PTZ group (n=3), which received the medial septum electrical stimulation for 30 min and PTZ injection (50 mg/kg, ip) (see below) 15 min after the septal stimulation started; 2) the 5 Hz + PTZ group (n=7), following the previously described protocol; and 3) the 5 Hz group (n=7), in which the septal electrical stimulation was delivered for 30 min without PTZ administration. The stimulation parameters consisted of a 30 min train of biphasic square-wave pulses, 10 Hz or 5 Hz, with a current of 150 µA and a pulse duration of 0.1 ms. Signal recordings were stopped 15 min after the electrical stimulation ended. Rats were submitted to the experimental protocol every 24 h for seven consecutive days.

### 2.5 PTZ-induced seizures

The rats were injected intraperitoneally with a high dose of PTZ (50 mg/kg, ip). Two experts evaluated the behavioral changes based on the following scheme: stage 1, motionless staring; stage 2, facial jerking; stage 3, neck jerks; stage 4, clonic seizure in a sitting position; stage 5, convulsions, including clonic and/or tonic-clonic seizures [48]. Stages 2 and 3 were considered focal seizures, while stages 4 and 5 were generalized seizures.

### 2.6 Brain extraction and histologic verification

Animals that finished the seven-day protocol were overdosed using an intraperitoneal injection of sodium pentobarbital (Pet’s Pharma). The rats were intracardially perfused with 0.9% NaCl solution followed by a 4% paraformaldehyde (PFA) solution (pH 7.4). The implanted setup was carefully removed. Then, the extracted brains were post-fixed in fresh 4% PFA solution for 24 h at 4 °C. Over the following days, tissues were immersed in 20% and 30% sucrose solutions at 4 °C. Each specimen was carefully frozen using dry ice and stored at −72 °C. The placement of the electrodes was verified by Nissl staining [49].

### 2.7 Signal analysis

Medial septum and left dorsal hippocampus signals were simultaneously recorded and analyzed in each rat by using the same time window for both structures. Signal analysis was performed offline. One 5 s epoch (digitized at 500 Hz) was chosen every day from each rat at three different time points: 1) baseline activity prior to the electrical stimulation, 2) during septal electrical stimulation prior to PTZ injection (specifically, 1 to 3 min before administering the pro-convulsive), and 3) during the activity preceding the onset of focal or generalized seizures (specifically, 30 to 60 s before the ictal period). When choosing the epochs, we avoided any abnormal artifact seen during the recording (e.g., signal saturation, spikes, noise due to animal movement or external factors). All analyses were conducted using Python 3.12, specifically the SciPy library and its signal processing functions.

For spectral analysis, the signals were digitized and sampled at 500 Hz and Notch-filtered at 60 Hz to remove line noise. The power spectral density was estimated using Welch’s method, which segments the signal length into a 1024-point window. The discrete Fourier transform was computed for each smoothed segment to estimate the spectral power. The periodograms from each window were averaged to create a more stable estimate of the power spectral density. The spectral analyses were performed for each structure, and frequency bands were defined as follows: delta (0-4 Hz), low-theta (4-8 Hz), high-theta (8-12 Hz), alpha (12-18 Hz), beta (18-30 Hz), slow-gamma (30-45 Hz), fast-gamma (45-60 Hz), and high-frequency oscillations (80-130 Hz). The relative power for each frequency band was calculated based on their area under the curve values.

### 2.8 Statistical analysis

The relative power data were normalized, and a repeated-measures ANOVA was used to compare daily septal and hippocampal activity at three different moments: baseline before electrical stimulation; during electrical stimulation prior to PTZ; and during pre-ictal activity. Next, we applied the Bonferroni *post hoc* test to determine differences among groups. For the group that did not receive the pro-convulsive injection, the analysis was done using paired *t*-tests. Values are expressed as mean ± S.E.M. In all statistical comparisons, significance was assumed at the level of *p* < 0.05.

Pearson correlation analysis was performed to assess the relationship between power values across conditions in both cerebral structures. Values are expressed as the mean. Statistical significance was determined using false discovery rate correction for multiple comparisons.

## 3. RESULTS

### 3.1 Histological analysis

None of the animals from the 10 Hz protocol were perfused since they died a few minutes after the pro-convulsive was injected. Additionally, we observed that the electrode tips were correctly implanted in both cerebral structures of the 5 Hz-stimulated animals (Fig. 1C and 1D).

### 3.2 Effects of medial septum electrical stimulation on PTZ-induced seizures

Animals were not allowed to sleep at any time. A 25 min visual examination of behavioral changes was performed once PTZ was administered. All the animals that received 10 Hz medial septum electrical stimulation had tonic-clonic seizures and died 5 to 10 min after the first injection. On the other hand, the seven rats from the 5 Hz group survived the whole week and presented focal seizures (FS) or generalized seizures (GS) as follows: none on day 1; 3 FS and 2 GS on days 2 and 3; 4 FS and 3 GS on day 4; 5 FS and 2 GS on day 5; 2 FS and 5 GS on day 6; and 7 FS on day 7. As seen, all the animals exhibited FS or GS, or no ictal activity throughtout the seven days. None of the GS lasted more than 90 s. Considering the 38 PTZ-induced convulsions, 63% were FS and 37% were GS.

Interestingly, we noticed medial septum spike-wave-like discharges (10 to 15 min after the electrical stimulation began) in animals in which generalization was not achieved. This protective effect was not observed prior to GS.

### 3.3 Local field potential power changes during 5 Hz stimulation and seizure induction

The evaluation focused on recordings in which seizures were induced (38/49). None of the analyses included the observed spike-wave-like discharges at the medial septum or its equivalent in the dorsal hippocampus.

During the 5 Hz stimulation, no significant changes were reported in the medial septum when any type of seizure was triggered (baseline vs electrical stimulation; Fig. 2). Before focal seizure onset, a significant increase in low-theta oscillations (baseline vs 1 min pre-ictal activity, *F_(2_*_,8)_ = 6.71, *p* < 0.05) was observed (Fig. 2B). Significances were achieved prior generalized seizures in both theta bands (low: *F*_(2,12)_ = 6.28, *p* < 0.01; high: *F*_(2,12)_ = 4.98, *p* < 0.05) (Fig. 2B and 2C).

**Figure 2.**
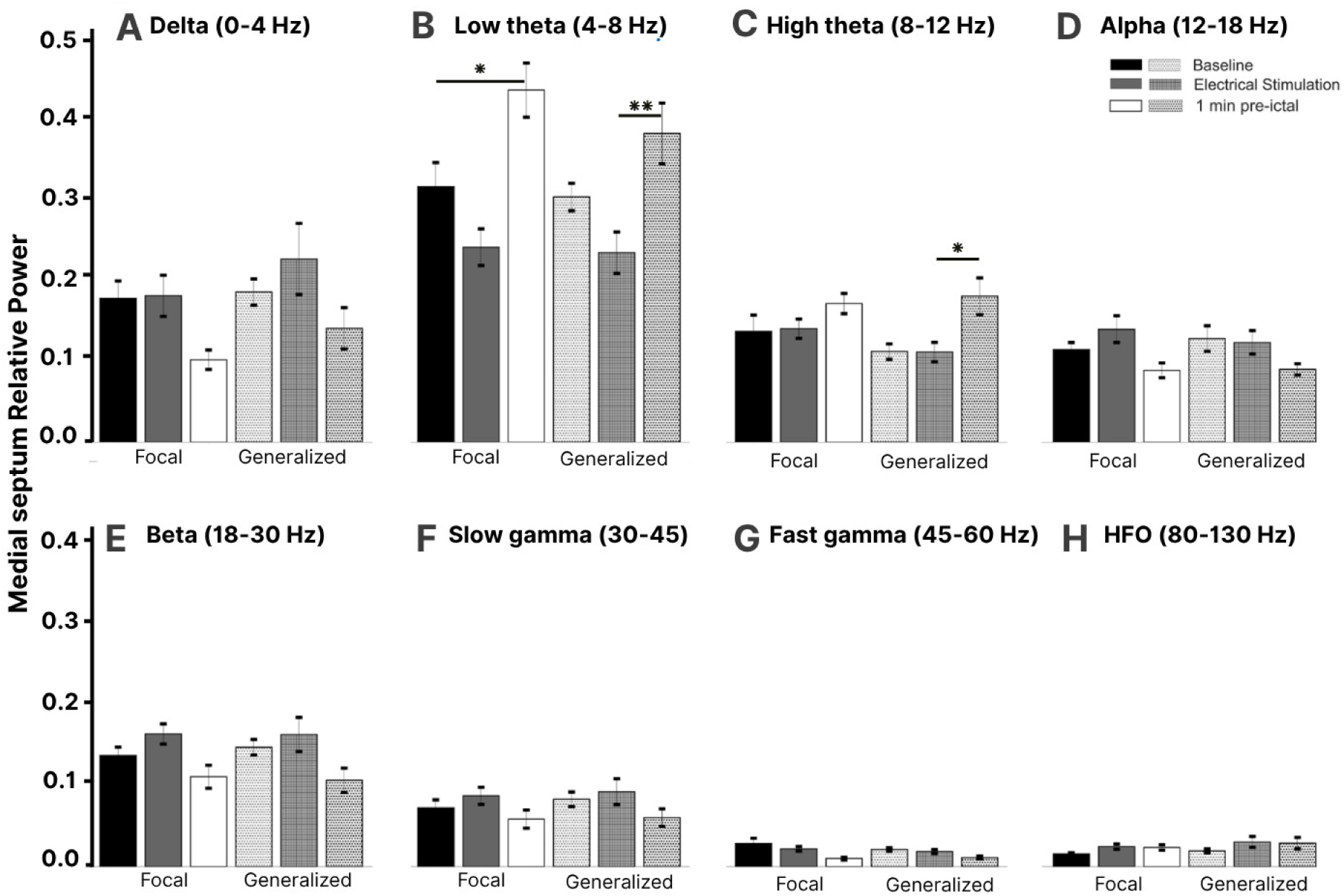
Mean relative power of the medial septum frequency bands during PTZ-induced seizures. Eight frequency bands (0 to 130 Hz) were analyzed **(A to H)** at three time points: 1) baseline prior to the electrical stimulation, 2) during septal electrical stimulation prior to PTZ (50 mg/ kg; ip) injection, and 3) during the activity preceding the onset of focal or generalized seizures. Before focal seizure onset, a significant increase in low theta oscillations was observed **(B)**. Significances were also achieved prior to generalized seizures in both theta bands **(B, C)**. Statistical comparisons between oscillations were performed using repeated-measures ANOVA followed by post hoc Bonferroni. \**p* < 0.05; \*\**p* < 0.01. HFO: High frequency oscillations.

The hippocampal focal seizure power analyses only showed significant differences in the slow-gamma oscillations (baseline vs electrical stimulation, *F*_(2,8)_ = 4.97, *p* < 0.01; Fig. 3F). Regarding the generalized seizures, significant decreases were observed in beta oscillations while comparing baseline vs electrical stimulation (*F*_(2,12)_ = 13.68, *p* < 0.05) and 1 min pre-ictal activity (*F*_(2,12)_ = 3.8, *p* < 0.05) (Fig. 3E).

**Figure 3.**
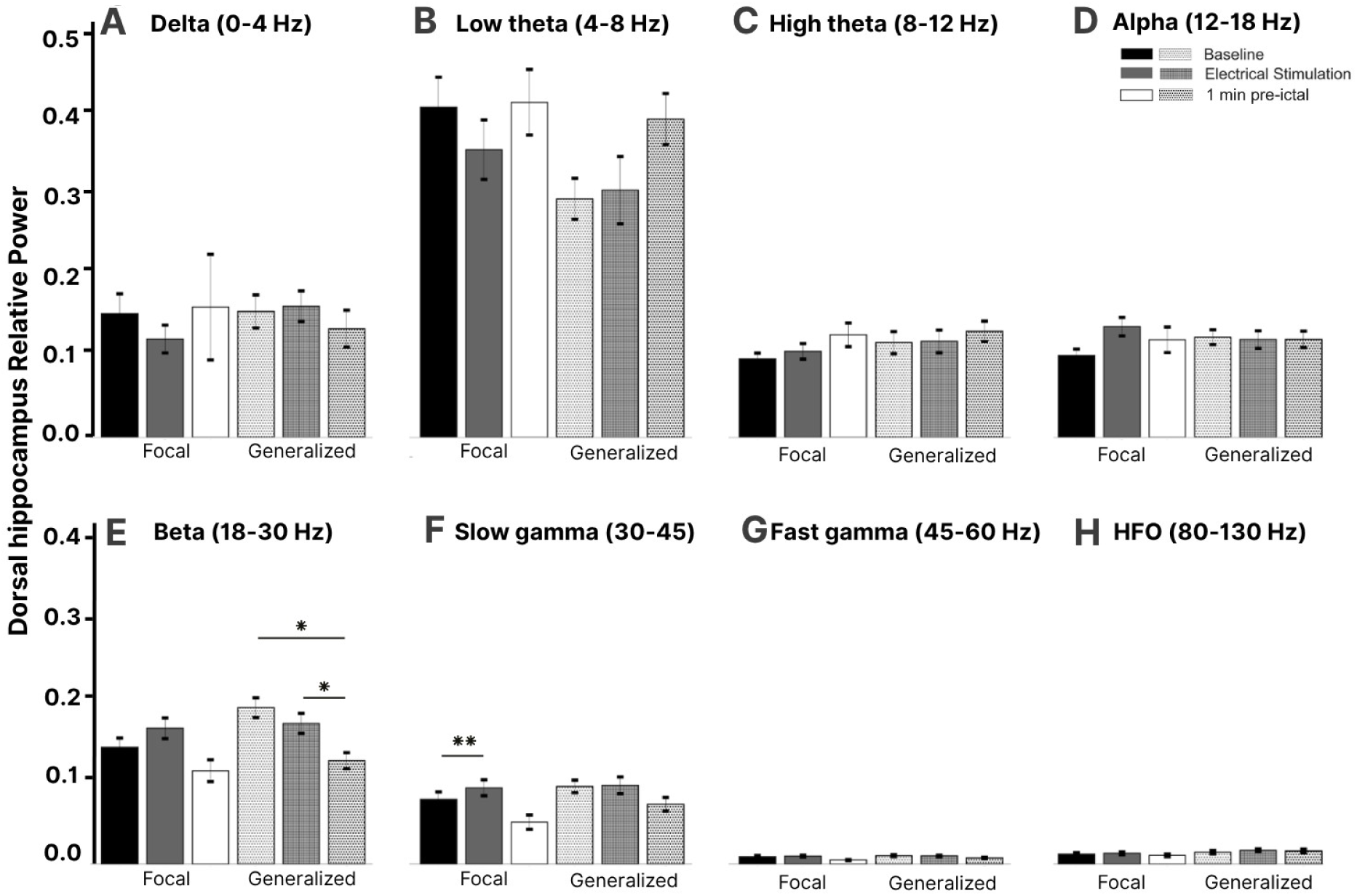
Mean relative power of the dorsal hippocampus frequency bands during PTZ-induced seizures. Delta (0 to 4 Hz) to high-frequency oscillations (HFO; 80 to 130 Hz) were evaluated **(A to H)**. The signal was analyzed at baseline prior to the electrical stimulation, during septal electrical stimulation (1 to 3 min before high-dose administration of the pro-convulsive), and 30 to 60 s prior to the ictal period. The focal seizure power analyses showed significant differences in the slow gamma oscillations **(F)**. Regarding generalized seizures, significant decreases were observed in beta oscillations when comparing baseline vs electrical stimulation and 1 min pre-ictal activity **(E)**. Statistical comparisons between bands were performed using repeated-measures ANOVA followed by post hoc Bonferroni. \**p* < 0.05; \*\**p* < 0.01.

Although no significant differences were reported after the correlation analyses (Fig. 4 and 5), electrical stimulation seemed to positively increase the correlation between structures in diverse bands such as high-theta, alpha, beta, slow-gamma, fast-gamma, and high-frequency oscillations (Fig. 4B and 5B). In the focal seizures analyses, a similar effect was detected (except for the high-theta band) (Fig. 4C). In contrast, the generalized seizures analyses showed values tending to zero or negative values in alpha, beta, and fast-gamma oscillations (Fig. 5C).

**Figure 4.**
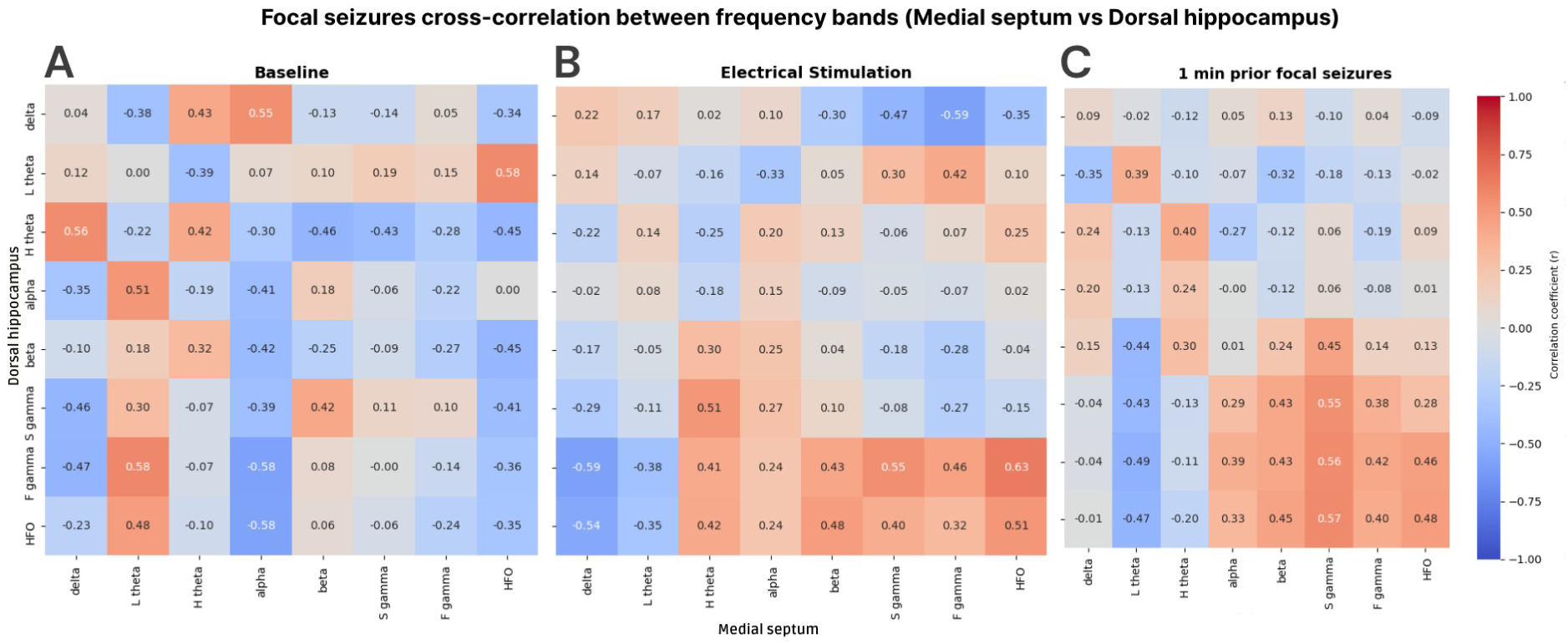
Correlations between medial septum (X-axis) and dorsal hippocampus (Y-axis) frequency bands prior to focal seizures. No significant differences were observed at any time point **(A to C)**. Nevertheless, electrical stimulation seems to positively increase the correlation between structures in high theta, alpha, beta, slow gamma, fast gamma, and high-frequency oscillations **(B)**. The 1 min prior focal seizures analysis shows a similar effect (except for high theta) **(C)**. Each cell represents the Pearson correlation coefficient (r). The colors indicate the magnitude and direction of the correlation: red for positive and blue for negative correlations; darker colors show stronger correlations. Statistical analysis was performed using multiple comparison correction by the False Discovery Rate (FDR) method. L: low; H: high; S: slow; F: fast; HFO: high-frequency oscillations.

**Figure 5.**
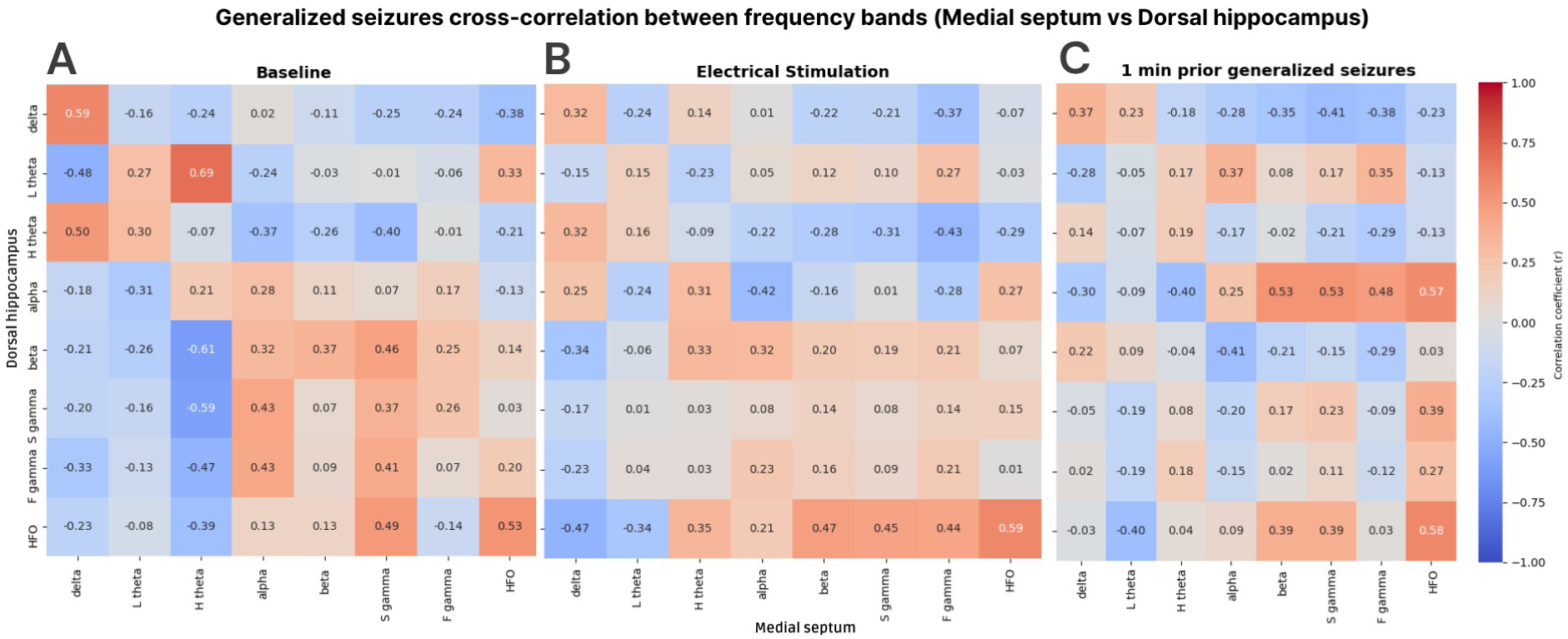
Correlations between medial septum (X-axis) and dorsal hippocampus (Y-axis) frequency bands prior to generalized seizures. Although no significant differences were observed at any time point **(A to C)**, electrical stimulation seems to positively increase the correlation between structures in high theta, alpha, beta, slow gamma, fast gamma, and high-frequency oscillations **(B)**. The 1 min prior generalized seizures analysis shows values tending to zero or negative values in alpha, beta, and fast gamma oscillations **(C)**. Each cell represents the Pearson correlation coefficient (r). The colors indicate the magnitude and direction of the correlation: red for positive and blue for negative correlations; darker colors show stronger correlations. Statistical analysis was performed using multiple comparison correction by the False Discovery Rate (FDR) method. L: low; H: high; S: slow; F: fast; HFO: high-frequency oscillations.

### 3.4 Local field potential power changes during 5 Hz stimulation in control rats

Although each rat was recorded and stimulated for seven consecutive days, here we only present the comparison of power values between days 1 and 7 from both evaluated structures (baseline vs electrical stimulation; Fig. 6 and 7). Animals were not allowed to sleep and none of them exhibited behavioral changes during the experimental protocol. However, during electrical stimulation, septal and hippocampal spike-wave-like discharges were seen in all rats (Fig. 9A). This phenomenon occurred 10 to 15 min after the 5 Hz stimulation began. The number of times these discharges appeared in each rat varied throughout the week (data not shown).

**Figure 6.**
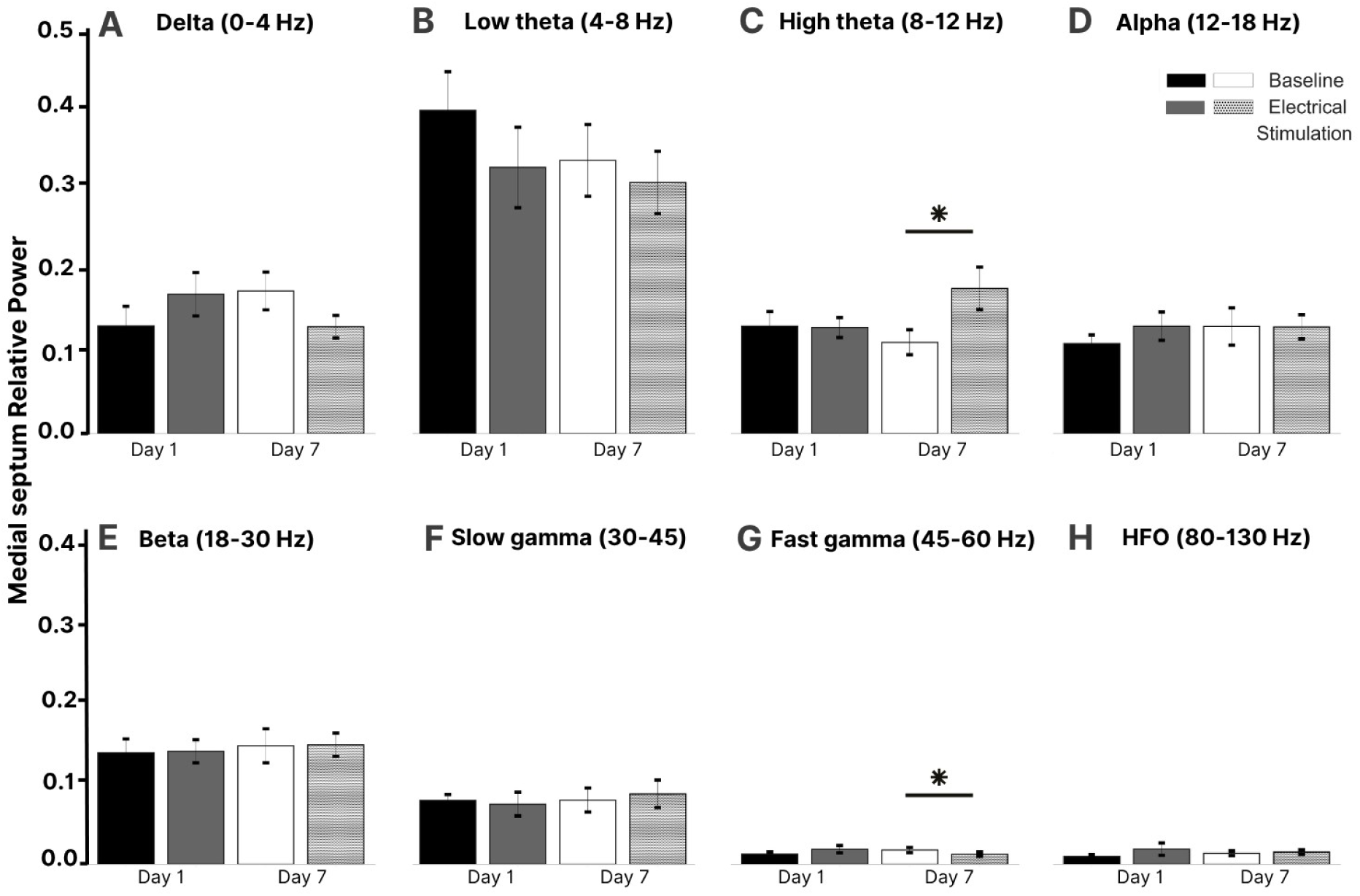
Mean relative power of the medial septum frequency bands in control rats. Eight frequency bands (0 to 130 Hz) were analyzed **(A to H)** at two time points: baseline and during the electrical stimulation on the first and last days of the experiment. On day 1, no significant differences were observed **(first two bars A to H)**. By the end of the week, significant differences were observed in high theta **(C)** and fast gamma oscillations **(G)**. Statistical comparisons between bands were performed using paired t-tests. \**p* < 0.05. HFO: high-frequency oscillations.

**Figure 7.**
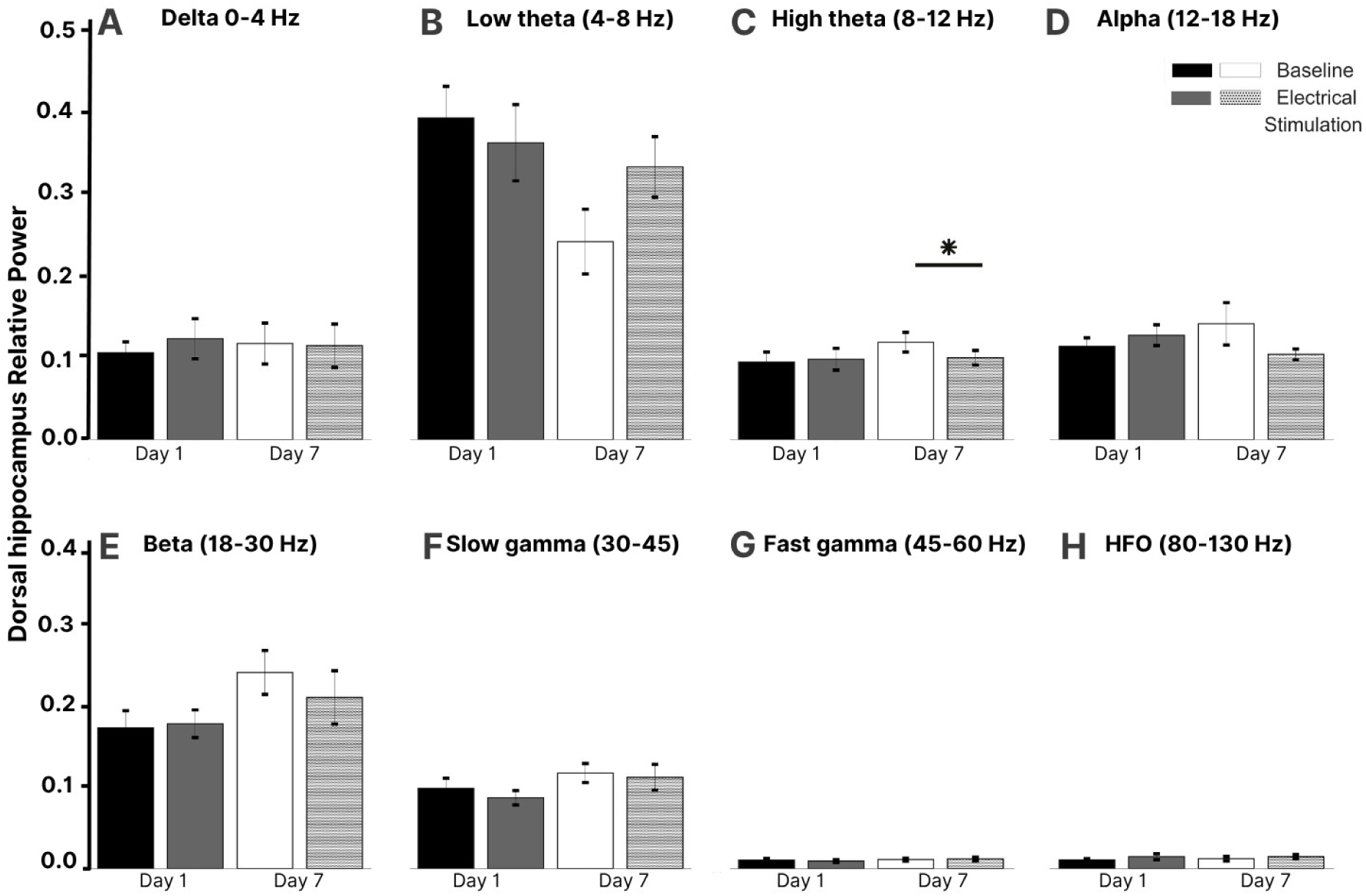
Mean relative power of the dorsal hippocampus frequency bands (0 to 130 Hz) in control rats. Delta (0 to 4 Hz) to high-frequency oscillations (HFO; 80 to 130 Hz) were evaluated **(A to H)**. During the first day of stimulation (day 1), no significant differences were observed **(first two bars A to H)**. On day 7, significance was only achieved in the high theta band **(C)**. Statistical comparisons between bands were performed using paired t-tests. \**p* < 0.05.

On day 1, no significant differences were observed (Fig. 6 and 7). By the end of the week, significant differences were observed in high-theta (*p* < 0.05; Fig. 6C) and fast-gamma oscillations (*p* < 0.05; Fig. 6G) in the medial septum. In the dorsal hippocampus, significance was only achieved in the high-theta band (*p* < 0.05; Fig. 7C). None of the analyses included the previously mentioned spike-wave discharges.

The correlation analysis showed no significant differences on day 1 (Fig. 8A and 8B). However, after seven days of 5 Hz septal stimulation, a significant positive correlation was observed in the baseline alpha band (r = 0.97, p_FDR = 0.027) (Fig. 8C). Moreover, during the electrical stimulation, a significant positive correlation was detected in fast-gamma oscillations (r = 0.98, p_FDR = 0.01) (Fig. 8D). This suggests that a daily 30 min septal stimulation can alter the circuitry dynamics.

**Figure 8.**
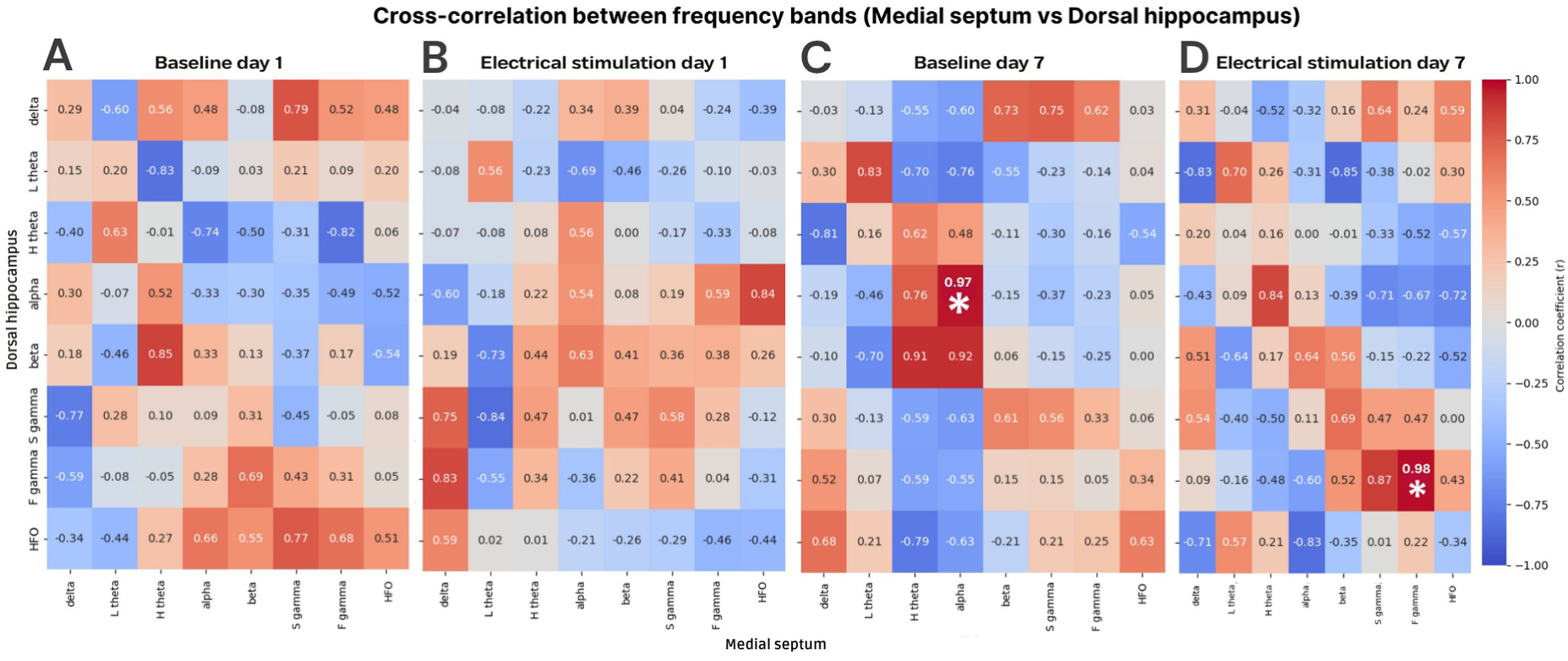
Correlations between medial septum (X-axis) and dorsal hippocampus (Y-axis) frequency bands in control rats. No significant differences were found on day 1 **(A, B)**. After seven days of electrical stimulation, a significant positive correlation was observed in the baseline alpha band **(C)**. Moreover, during the electrical stimulation, a significant positive correlation was detected in the fast gamma oscillations **(D)**. Each cell represents the Pearson correlation coefficient (r). The colors indicate the magnitude and direction of the correlation: red for positive and blue for negative correlations; darker colors show stronger correlations. Statistical analysis was performed using multiple comparison correction by the False Discovery Rate (FDR) method. \**p* < 0.05. L: low; H: high; S: slow; F: fast; HFO: high-frequency oscillations.

**Figure 9.**
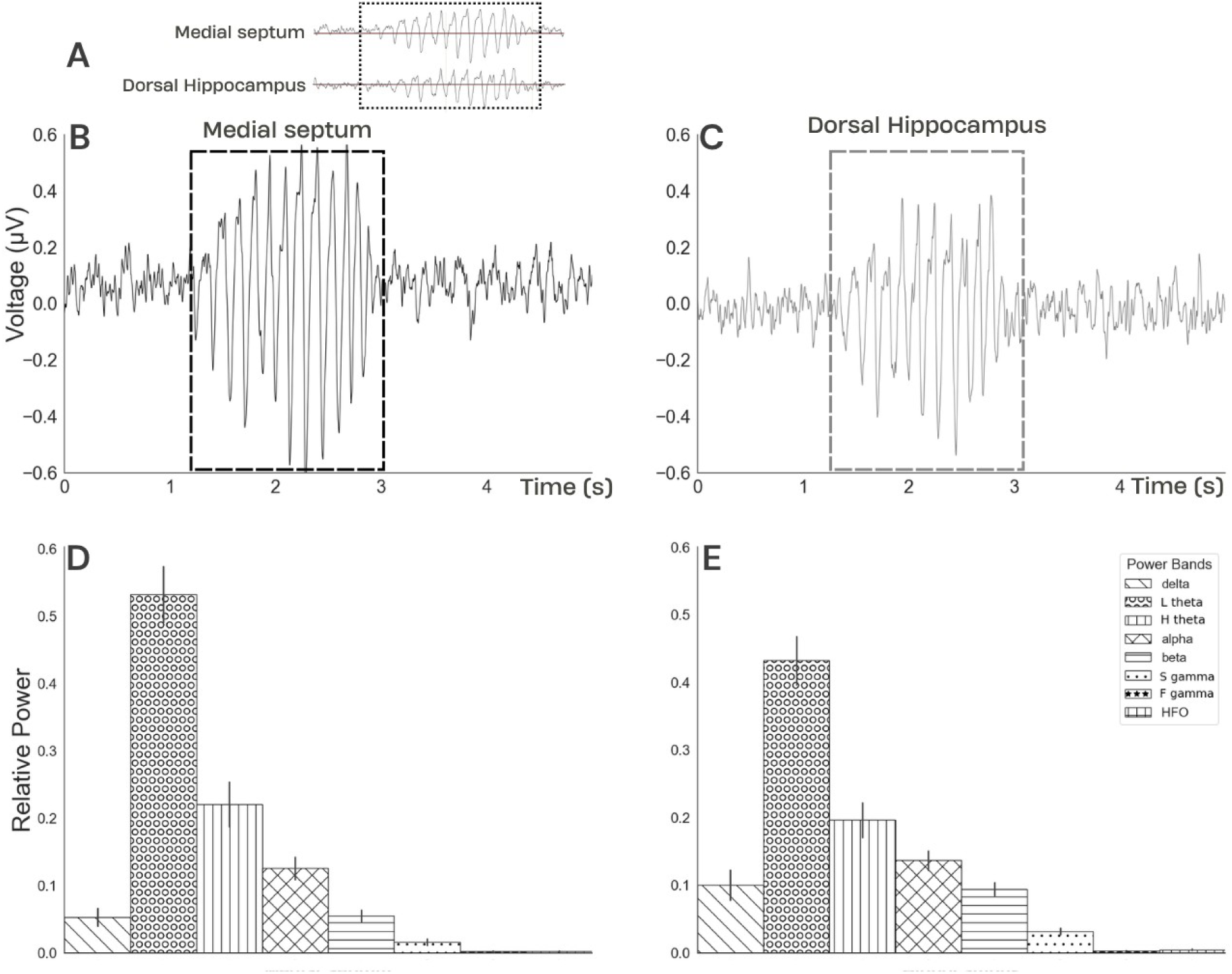
Mean relative power of the spike-wave-like discharges in control rats. Septal and hippocampal spike-wave-like discharges were seen in all the stimulated rats at 5 Hz. Twenty-six randomly chosen clusters were analyzed. The spike-wave-like discharges were simultaneously recorded in the medial septum and left dorsal hippocampus **(A)**. Their duration was no longer than 2 s, and their frequency was predominantly between 4 and 12 Hz **(B, C)**. Theta oscillations cover a high percentage of the activity described for both the medial septum **(D)** and dorsal hippocampus **(E)**.

### 3.5 Spike-wave-like discharges analysis

The spike-wave-like discharges were simultaneously recorded in both evaluated structures of the electrically stimulated control rats (Fig. 9A). Moreover, the local field potential traces demonstrate that they exhibited a high similarity to septal or hippocampal activity throughout the week (data not shown). The spike-wave-like discharges were easily identified due to the spikes with an ascending phase (at least twice the amplitude of the background), followed by an abrupt descending phase. Their duration was no longer than 2 s, and their frequency was predominantly between 4 and 12 Hz (Fig. 9B and 9C). During these events, we did not notice the animals in stationary postures or with evident decreased activity.

For the analysis, we randomly chose 26 spike-wave-like discharges (daily, and across all animals). Then, the spike-wave-like discharges from both structures were analyzed using 2 s epochs to perform the corresponding power analyses. Theta oscillations covered a high percentage of the activity described for both the medial septum (near 75%; Fig. 9D) and the dorsal hippocampus (near 65%; Fig. 9E).

## 4. DISCUSSION

This study was designed to assess the effect of low-frequency deep brain stimulation of the medial septum to inhibit PTZ-induced seizures in rats. The main findings were: 1) The 10 Hz medial septum stimulation for 30 min had no effect in preventing severe generalized seizures and subsequent animal death. 2) The 5 Hz medial septum stimulation for 30 min increased the animals’ survival; throughout the week, on some days it induced anticonvulsive effects, while on other days it had a protective effect against generalization. 3) The presence of medial septum spike-wave-like discharges when no generalization was achieved. 4) The 5 Hz medial septum stimulation induced repetitive septal and hippocampal spike-wave-like discharges in control rats. 5) Spike-wave-like discharges are mainly integrated by theta oscillations.

Deep brain stimulation is an option when surgery has failed or is not appropriate for certain refractory epileptic patients [50]. Clinical and preclinical studies have shown its efficacy in reducing seizure frequency [51,52]. Experimental models have even compared the anticonvulsive effects when applying low- or high-frequency electrical stimulation to specific brain regions [14,53–58]. However, evident conflicting results have been reported among these studies.

The anti-ictogenic effect that we observed is consistent with previous evidence that deep brain electrical stimulation of the medial septum decreases seizure occurrence [35,38,40]. In this study, we suggest that the site of stimulation (medial septum), the frequency of stimulation (5 Hz), and the train duration (30 min) may be key factors in protecting the animals against secondary generalization and death. Over the past decade, the medial septum has been proposed as a potential target for the treatment of drug-resistant temporal lobe epilepsy [16,17]. This is mainly attributed to its location along the midline, which eliminates the need for bilateral electrode implantation and its connectivity characterized by strong projections to the hippocampus and entorhinal cortex impacting theta frequency activity and seizure generalization [22,59–62]. Regarding the frequency of stimulation, diverse experimental studies have reported that low-frequency medial septum stimulation (ranging between 0.5 and 7.7 Hz) has more effective antiseizure effects than higher stimulation frequencies [35,38–40,63]. In some cases, this is caused by increased hippocampal theta band power [38,40,63] or optogenetic stimulation of specific medial septum neuronal populations [37,39,41,64]. Although research has demonstrated the beneficial effects of low-frequency medial septum stimulation on experimental seizures, little is known about the train duration. Nevertheless, studies on the effects of thalamic electrical stimulation have established that train duration and stimulation protocols are crucial factors in protecting animals against secondary generalization, *status epilepticus*, or even death as a consequence of hyperexcitability [58,65].

Additionally, our results showed that electrical stimulation alone induced a significant increase in the hippocampal power ratio of slow-gamma oscillations (30-45 Hz) only when no generalized seizures were achieved. Other significant changes were observed during the 30-min electrical stimulation protocol, but before focal or generalized seizure onset. Specifically, theta oscillations (4-12 Hz) increased in the medial septum, while beta oscillations (18-30 Hz) decreased in the dorsal hippocampus. Recently, Zhang et al. [63] reported that the decrease in septal gamma oscillations and increase in beta oscillations after deep brain stimulation of the septal area could be a potential mechanism for attenuating kainic acid-induced seizures. Other authors, while evaluating learning and memory, have reported the relevance of the medial septum during these processes due to the cross-frequency coupling between gamma and theta oscillations in this brain structure [66,67]. Regarding theta oscillations, an uncoupling event between septal activity and hippocampal theta rhythm seems to play a crucial role in epileptogenesis [30,68,69]. Local field potential recordings from epileptic rats have shown a significant decrease in hippocampal theta power and amplitude [28,70]. Moreover, lesions of the medial septum in rodents also attenuate them [71,72]. Diverse studies have described that these oscillations are mainly regulated by septal cholinergic input into the hippocampal formation and play a crucial role in seizure control [63,73,74]. However, while it is evident that low-frequency medial septum stimulation is correlated with anticonvulsive effects, research has yet to determine the precise mechanism that makes it a promising anti-ictogenic option.

Studies from our laboratory and others suggest that low-frequency stimulation of the medial septum may regulate oscillatory activity in proximal and distal connected regions such as the hippocampus. Interestingly, in this study we show for the first time that this type of stimulation also induces septal and hippocampal spike-wave-like discharges, mainly integrated by theta oscillations in both cases. This phenomenon represents a major electrophysiological characteristic of absence epilepsy. Common models for experimental studies are GAERS (Genetic Absence Epilepsy rat from Strasbourg), WAG/Rij (Wistar Absence Glaxo from Rijswik), and myelin mutant taiep rats (characterized by tremor, ataxia, immobility episodes, epilepsy, and paralysis) [75,76]. The generation of spike-wave discharges has usually been associated with the cortico-thalamo-cortical system [77,78]. However, although no spike-wave discharges have been recorded from the hippocampal formation, numerous studies have not ruled out the involvement of limbic structures in such events [79–81]. Specifically, Papp et al. [81] argue that alterations in cortical and hippocampal GABAergic interneurons may damage the inhibitory mechanisms and contribute to the emergence of absence epileptic spike-wave discharges. On the other hand, it is worth noting that GAERS and WAG/Rij rats are also known to be resistant to secondary generalization of focal limbic seizures in response to kindling protocols. An effect that has been related to augmented hippocampal GABAergic activity [82–86]. To date, there is no evidence of spike-wave discharges in the medial septum of any rat strain. Regarding the relationship between deep brain stimulation and such discharges, we found a couple of reports where depending on the frequency stimulation protocol, spike-wave discharges were induced, maintained, disrupted, or decreased [58,87].

Given that neurostimulation for epilepsy focuses on either the seizure onset zone or a relevant network node affecting that area, our findings add further evidence to the notion that medial septum low-frequency stimulation may represent an effective treatment for drug-resistant patients with focal seizures. In addition to the anticonvulsive or generalized-suppression effect, we report for the first time that the 5 Hz protocol induced septal and hippocampal spike-wave-like discharges, suggesting that stimulation of the medial septum could induce theta activity via these clusters. Future goals include the adaptation of this protocol to rat models of recurrent severe seizures and drug-resistant epilepsy.

## 5. LIMITATIONS

It is important to mention some considerations about our study. First, our protocol only evaluated the antiseizure effect of the electrical stimulation. To evaluate how this strategy can improve outcomes, we need to adapt the protocol to rat models of recurrent severe seizures and drug-resistant temporal lobe epilepsy. Results from these studies will significantly help translatability to clinical settings. Second, we did not evaluate the long-term benefits of stimulation nor analyze changes in cell death, neuroinflammation or neurotransmission. *Ex vivo* anatomical magnetic resonance imaging studies were not possible due to tissue damage caused by the electrodes and by implant removal. Overcoming these challenges will add valuable information to our data.

## 6. CONCLUSION

We demonstrate, in a rodent model widely used in the search for new antiseizure drugs, that the medial septum and theta oscillations represent relevant targets for reducing seizure severity and, in some cases, prevent seizure onset. Moreover, the results show, for the first time, that 5 Hz medial septum stimulation induces spike-wave-like discharges in control rats and that this phenomenon was also observed in animals without PTZ-induced generalized seizures. While this study is preclinical, our findings underscore the potential of using low-frequency medial septum stimulation for future clinical applications. Future experiments are crucial to determine the mechanism of action of such a strategy and understand/prevent possible secondary effects after long-lasting stimulation protocols.

## Statements & Declarations

### Ethics approval and consent

This research was approved by the Institute of Neurobiology Ethics Animal Care Committee (protocol 105A).

### Author Contributions

AG-C, VM-M, and HL-M contributed to the conception, methodology, and design of the study. AG-C, SA-A, VM-M, and HL-M performed local field potentials acquisition and histological procedures. AG-C, VM-M, and HL-M performed and supervised statistical analyses. AG-C, VM-M, and HL-M wrote, reviewed, and edited the first manuscript. All authors reviewed the manuscript and approved the submitted version.

### CRediT authorship contribution statement

**Alejandra Garay-Cortes:** Data curation, Formal analysis, Investigation, Methodology, Writing – original draft. **Salvador Almazan-Alvarado:** Data curation, Methodology. **Victor Magdaleno-Madrigal:** Data curation, Formal analysis, Methodology, Resources, Supervision, Validation, Visualization, Writing – review and editing. **Hiram Luna-Munguia:** Conceptualization, Data curation, Formal analysis, Funding acquisition, Investigation, Methodology, Project administration, Resources, Supervision, Validation, Visualization, Writing – review and editing.

### Data availability

All data generated and analyzed during this study will be made available by the authors, without undue reservation.

### Ethics approval

The animal study was reviewed and approved by the Ethics Committee of the Institute of Neurobiology at the Universidad Nacional Autonoma de Mexico.

## Acknowledgments

Alejandra Garay-Cortes is a doctoral student of the Programa de Doctorado en Ciencias Biomedicas at Universidad Nacional Autonoma de Mexico (UNAM) and has received a fellowship (CVU 1005958) from Secretaria de Ciencias, Humanidades, Tecnologia e Innovacion (SECIHTI, formerly CONAHCYT). We thank Mirelta Regalado, Juan Ortiz-Retana, Ericka de los Rios, Nydia Hernandez, Lourdes Palma, Moises Mendoza, and Leopoldo Gonzalez for their technical assistance. We also thank the personnel at the Animal Facility: Martin Garcia, Alejandra Castilla, Maria Carbajo-Mata, and Maria Eugenia Ramos. Jessica Gonzalez-Norris for proof-reading and editing our manuscript. MRI preclinical scanner facilities were provided by the National Laboratory of Magnetic Resonance (LANIREM), which receives support from SECIHTI and UNAM. Computing infrastructure was partially provided by the National Laboratory for Advanced Scientific Visualization (LAVIS).

## Declaration of generative AI use

The authors declare no use of generative AI in the manuscript preparation process.

